# Evaluation of Natural and Botanical Medicines for Activity against Growing and Non-growing Forms of *B. burgdorferi*

**DOI:** 10.1101/652057

**Authors:** Jie Feng, Jacob Leone, Sunjya Schweig, Ying Zhang

## Abstract

Lyme disease is the most common vector-borne disease in the US. Although the current recommended Lyme antibiotic treatment can cure the majority of Lyme disease patients, about 10-20% patients continue to suffer from persisting symptoms. There have been various anecdotal reports on the use of herbal extracts for treating patients with persisting symptoms with varying degree of improvements. However, it is unclear whether the effect of the herb products is due to their direct antimicrobial activity or their effect on host immune system. In the present study, we investigated the antimicrobial effects of 12 commonly used botanical medicines and 3 other natural antimicrobial agents for potential anti-*Borrelia burgdorferi* activity in vitro. Primary criteria for selecting compounds for the present study included agents that had shown significant anti-borrelial effects in previous studies, have favorable safety profiles, and can be absorbed systemically. Among them, 9 natural product extracts at 1% were found to have good activity against the stationary phase *B. burgdorferi* culture compared to the control antibiotics doxycycline and cefuroxime. These active herbs include *Cryptolepis sanguinolenta, Juglans nigra* (Black walnut), *Polygonum cuspidatum* (Japanese knotweed), *Artemesia annua* (Sweet wormwood), *Uncaria tomentosa* (Cat’s claw), *Cistus incanus*, and *Scutellaria baicalensis* (Chinese skullcap). In contrast, *Stevia rebaudiana, Andrographis paniculata*, Grapefruit seed extract, colloidal silver, monolaurin, and antimicrobial peptide LL37 had little or no activity against stationary phase *B. burgdorferi*. The minimum inhibitory concentration (MIC) values of *Artemesia annua, Juglans nigra*, and *Uncaria tomentosa* were quite high for growing *B. burgdorferi*, despite their strong activity against the non-growing stationary phase *B. burgdorferi* cells. On the other hand, the top two active herbs, *Cryptolepis sanguinolenta* and *Polygonum cuspidatum*, showed strong activity against both growing *B. burgdorferi* (MIC=0.03%-0.06% and 0.25%-0.5% respectively) and non-growing stationary phase *B. burgdorferi*. In subculture studies, only 1% *Cryptolepis sanguinolenta* extract caused complete eradication, while current Lyme antibiotics doxycycline and cefuroxime and other active herbs including *Polygonum cuspidatum, Artemesia annua, Juglans nigra* and *Uncaria tomentosa* could not eradicate *B. burgdorferi* stationary phase cells as many spirochetes were visible after 21-day subculture. Further studies are needed to identify the active ingredients of the effective herbs and evaluate their combinations for more effective eradication of *B. burgdorferi* in vitro and in vivo. The implications of these findings for more effective treatment of persistent Lyme disease are discussed.

## Introduction

Lyme disease, caused by *Borrelia burgdorferi*, and multiple closely related *Borrelia* species, is the most common vector-borne human disease in the Northern Hemisphere (1, 2). About 300,000 new cases are reported in the United States annually (3, 4). Tick-borne infections are on the rise in the USA and Europe due to a host of different factors including climate change (5–7), and disruption of predator density in suburban areas (8). Recent studies on tick prevalence and pathogen load have identified new geographical areas where vector ticks are present (9), as well as novel tick-borne pathogens present in areas where they had not previously been identified (such as *B. miyamotoi* in Northern California)(10).

Lyme disease can affect many different body systems and organs (11). While many patients recover fully with early antibiotic therapy, at least 10-20% of patients experience persistent symptoms following the conventionally recommended course of 2-4 weeks of antibiotics (12–14), and a recent retrospective analysis documented 63% of patients experienced persistent symptoms after receiving antibiotic treatment for Lyme disease (15). Patients who experience persistent symptoms have significant and ongoing disability (16, 17) and increased health care costs and utilization (15). *B. burgdorferi* can evade the immune system response (18, 19) and multiple studies have shown that the bacteria is capable of persisting in diverse tissues across a variety of animal models despite aggressive and prolonged antibiotic therapy (20–23).

In addition to the mammalian studies noted above, *B. burgdorferi* persistence following antibiotic treatment has been demonstrated in human studies and case reports (24–27). Persistent Lyme borreliosis symptoms significantly affect quality of life (28, 29), therefore some physicians treat these patients with extended courses of antibiotics. However, this approach is controversial with one medical society guideline (30) advocating against retreating patients with persistent (> 6 months) symptoms and another medical society guideline (31) recommends individualized risk-benefit assessments and potential retreatment or longer duration treatment of patients with persistent symptoms. While antibiotic retreatment has been associated with improved clinical outcomes (31, 32) it is of vital importance that novel safe and effective treatments be identified for clinical use. Furthermore, traditional antibiotic therapy appears to be more effective against the actively dividing spirochete form, and it has been shown that *B. burgdorferi* can change morphology and form biofilm-like microcolonies consisting of stationary phase persister bacteria (33–35). Traditional antibiotics have poor activity against the atypical persister forms (round bodies, microcolonies, and biofilm) and we have previously worked to identify novel drugs and drug combinations that are effective (33, 35, 36). While Daptomycin and Dapsone have been identified as having significant effects against borrelia persister cells in vitro (35, 37) and in vivo in a murine model (34), their use in clinical practice can be limited by side effects (both), cost (daptomycin), parenteral administration (daptomycin) and poor CNS penetration (daptomycin) (38).

Importantly, botanical medicines have been shown to have in vitro antimicrobial activity against various morphologic forms of *B. burgdorferi*. Because there are a limited number of studies evaluating the effects of botanical medicine on *B. burgdorferi*, it is helpful to draw on clinical studies that have shown benefit using botanical medicines for other spirochetal infections and infections like mycobacterium that are known to form antibiotic tolerant persister cells (39). For example, *Andrographis* has been shown to effectively treat leptospirosis in Chinese clinical trials (40, 41) and improve clinical outcomes when combined with standard antituberculosis treatment for TB (42).

Botanical medicine has a long history of use, beginning almost 5000 years ago in Mesopotamia and over 3000 years of documented usage in China (43, 44). The safety of botanical medicines has been documented in various traditional systems of medicine such as Ayurvedic Medicine and Traditional Chinese Medicine over centuries. Recent retrospective and systematic reviews in the European Union and South America have concluded severe adverse events associated with Botanical Medicine usage were rare (45–47).

This study builds on previous studies that used our in vitro stationary phase persister model and SYBR Green I/propidium iodide (PI) assay to screen potential antimicrobial candidates. Having previously identified novel drugs and drug combinations from an FDA drug library (36), as well as selected botanicals in essential oil form that have anti-*B. burgdorferi* activity (48, 49), in the present study we investigated the effect of 12 botanical medicines and 3 other natural antimicrobial agents for potential anti-*B. burgdorferi* activity in vitro.

## Materials and Methods

### Strain, media, and culture techniques

*B. burgdorferi* strain B31 was cultured in BSK-H medium (HiMedia Laboratories Pvt. Ltd.) with 6% rabbit serum (Sigma-Aldrich, St. Louis, MO, USA). All culture medium was filter-sterilized by 0.2 µm filter. Cultures were incubated in sterile 50 ml conical tubes (BD Biosciences, CA, USA) in microaerophilic incubator (33°C, 5% CO_2_) without antibiotics.

### Botanical and natural medicines

A panel of natural product extracts: *Polygonum cuspidatum, Cryptolepis sanguinolenta, Artemisia annua, Juglans nigra, Uncaria tomentosa, Scutellaria baicalensis, Stevia rebaudiana, Cistus incanus, Andrographis paniculata* (Chuan Xin Lian), *Ashwagandha somnifera, Dipsacus fullonum rad*, grapefruit seed extract, LL37, monolaurin, colloidal silver and relevant solvent controls (see Table 2) were identified. The botanical medicines or natural products were chosen based on anecdotal clinical usage and preclinical data from the literature. Primary criteria for selecting compounds for the present study included agents that had shown significant anti-borrelial effects in previous studies, have favorable safety profiles and can be absorbed systemically. Additional criteria for selecting compounds included anecdotal reports from patients and/or providers, anti-biofilm effects and ability to cross the blood brain barrier.

**Table 1.** Activity of natural products against growing (MIC) and stationary phase *B. burgdorferi*.

| Natural products | MIC (%) <sup>a</sup> | Stationary phase residual viability (%) at different concentrations of herbs <sup>b</sup> |  |  | Subculture |  |
| --- | --- | --- | --- | --- | --- | --- |
|  |  | 1% | 0.5% | 0.25% | 1% | 0.5% |
| Drug free control |  | 94% |  |  | + |  |
| 5 µg/ml Doxycycline | 0.25 µg/mL | 74% |  |  | + |  |
| 5 µg/ml Cefuroxime | 0.13 µg/mL | 65% |  |  | + |  |
| 30% alcohol control | >2% | 79% | 80% | 95% | + | + |
| 60% alcohol control | 1%-2% | 77% | 76% | 94% | + | + |
| 90% alcohol control | 0.5%-1% | 75% | 79% | 91% | + | + |
| *Polygonum cuspidatum* 60% EE | **0.25%-0.5%** | **30%** | **41%** | **43%** | + | + |
| *Cryptolepis sanguinolenta* 60% EE | **0.03%-0.06%** | **46%** | **48%** | **46%** | - | +<sup>c</sup> |
| *Artemisia annua* 90% EE | 0.5%-1% | **43%** | **50%** | **49%** | + | + |
| *Juglans nigra* 60% EE | 0.5%-1% | **14%** | **36%** | **53%** | + | + |
| *Uncaria tomentosa* (inner bark) WE | 1%-2% | **49%** | **47%** | **54%** | + | + |
| *Polygonum cuspidatum* 90% EE | **0.25%-0.5%** | **21%** | **43%** | 61% | + | + |
| *Juglans nigra* 30% EE | 1%-2% | **33%** | **50%** | 62% | + | + |
| *Scutellaria baicalensis* | >2% | **59%** | 60% | 62% | + | + |
| *Cryptolepis sanguinolenta* 90% EE | **0.03%-0.06%** | **48%** | **47%** | 63% | ND | ND |
| *Juglans nigra* 90% EE | 0.5%-1% | **34%** | **56%** | 63% | ND | ND |
| *Cryptolepis sanguinolenta* 30% EE<sup>d</sup> | **0.06%-0.13%** | **59%** | **64%** | 63% | ND | ND |
| *Juglans nigra* fruc | 1%-2% | **52%** | **59%** | 66% | ND | ND |
| *Scutellaria baicalensis* 60% EE | **0.25%-0.5%** | 62% | 67% | 67% | ND | ND |
| *Scutellaria baicalensis* 90% EE | **0.25%-0.5%** | 72% | 74% | 75% | ND | ND |
| *Andrographis paniculata* 90% EE | 0.5%-1% | 74% | 75% | 75% | ND | ND |
| *Scutellaria baicalensis* 30% EE | **0.25%-0.5%** | 80% | 72% | 77% | ND | ND |
| *Cistus incanus* | 0.25%-0.05% | **29%** | 74% | 77% | ND | ND |
| *Andrographis paniculata* 30% EE | 1%-2% | 79% | 78% | 78% | ND | ND |
| Chuan Xin Lian | >2% | 89% | 86% | 85% | ND | ND |
| Citrosept <sup>TM</sup> | 1%-2% | 89% | 90% | 85% | ND | ND |
| *Polygonum cospidatum* 30% EE<sup>d</sup> | **0.25%-0.5%** | **34%** | 65% | 87% | ND | ND |
| Lauricidin <sup>TM</sup> | >2% | 88% | 86% | 87% | ND | ND |
| *Scutellaria barbata* | >2% | 58% | 60% | 88% | ND | ND |
| *Stevia rebaudiana* fol | >2% | 86% | 66% | 88% | ND | ND |
| *Andrographis paniculata* 60% EE | 1%-2% | 76% | 77% | 88% | ND | ND |
| *Dipsacus fullonum* rad | >2% | 84% | 90% | 89% | ND | ND |
| LL37 antimicrobial peptide | >2% | 91% | 91% | 89% | ND | ND |
| *Uncaria tomentosa* | >2% | 68% | 90% | 91% | ND | ND |
| *Ashwagandha somnifera* 90% EE | 0.5%-1% | 76% | 76% | 92% | ND | ND |
| *Ashwagandha somnifera* 60% EE | 0.5%-1% | 79% | 81% | 92% | ND | ND |
| Colloidal silver (Argentyn <sup>TM</sup> ) | >2% | 88% | 85% | 92% | ND | ND |
| *Ashwagandha somnifera* 30% EE | 0.5%-1% | 94% | 94% | 93% | ND | ND |
| Citrosept <sup>TM</sup> | 1%-2% | 98% | 99% | 95% | ND | ND |
| Grapefruit seed extract | Citrus paradisi | 78% | 81% | 94% | ND | ND |

<sup>a</sup> The standard microdilution method was used to determine the minimum inhibitory concentration (MIC). The MICs below 0.5% are shown in bold.
<sup>b</sup> A 7-day old *B. burgdorferi* stationary phase culture was treated with natural product extracts or control drugs for 7 days. Bold type indicates the samples that had better activity compared with doxycycline or cefuroxime controls. Residual viable *B. burgdorferi* was calculated according to the regression equation and ratios of Green/Red fluorescence obtained by SYBR Green I/PI assay.
<sup>c</sup> One of triplicate subculture samples grew up, and the other two samples did not grow back. Abbreviations: EE: ethanol extract; WE: water extract.

**Table 2:** Botanical and natural medicine sources, validation, and testing.

| **Natural Product** | **Source** | **Validation/ID** | **Contamination** | **Details** |
| --- | --- | --- | --- | --- |
| *Citrus x paradisi* | Cintamani, Poland (Citrosept™) | Cintamani, Poland | <1ppm for Benzalkonium chloride, Triclosan, Benzoic Acid | Organic grapefruit seed extract |
| *Stevia rebaudiana* | Sonoma County Herb Exchange (Cultivated) | Organoleptic, KW Botanicals | N/A | 25% ETOH extract by KW Botanicals |
| *Juglans nigra* | Pacific Botanicals (Wild harvested) | Organoleptic, KW Botanicals | N/A | 45% ETOH extract of husk/hulls by KW Botanicals |
| *Dipsacus fullonum* | Friend's of the Trees (wild harvested, Washington State) | DNA Species Identification, NSF International | N/A | 40% ETOH by KW Botanicals (Inadvertently co-mingled with D. asper sample prior to testing) |
| *Dipsacus asper* | KW Botanicals (Wild harvested, California) | DNA Species Identification, NSF International | N/A | 40% ETOH by KW Botanicals (Inadvertently co-mingled with D. fullonum sample prior to testing) |
| *Uncaria tomentosa* | Mountain Rose Herbs (Wild harvested) | DNA Species Identification, Christopher Hobbs, Ph.D. | negative testing for aerobic plate count, e. coli, coliform, salmonella, yeast & mold | 50% ETOH by KW Botanicals |
| *Artemisia annua* | Heron Botanicals (Organic cultivation) | American Herbal Pharmacopoeia (Scotts Valley, CA), Organoleptic, Heron Botanicals<br>Confirmed 0.11% Artemisinin content, The Institute for Food Safety and Defense | negative testing for aerobic plate count and yeast & mold | 30, 60, 90% ETOH by Heron Botanicals |
| *Withania somnifera* | Heron Botanicals (Organic cultivation) | HPTLC, The Institute for Food Safety and Defense<br>Organoleptic, Heron Botanicals | negative testing for Pb, Cd, Hg, As, aerobic plate count and yeast & mold | 30, 60, 90% ETOH by Heron Botanicals |
| *Juglans nigra* | Heron Botanicals (Wild harvested, New York) | Organoleptic, Heron Botanicals | positive aerobic plate count: 960 CFU/ml (acceptable limit 1,000 CFU/ml)<br>negative testing for Pb, Cd, Hg, As, and yeast & mold | 30, 60, 90% ETOH by Heron Botanicals |
| *Andrographis paniculata* | Heron Botanicals<br>(Organic cultivation, China) | Organoleptic, Heron Botanicals | negative testing for pesticides, sulfur dioxide, aerobic plate count and yeast & mold | 30, 60, 90% ETOH by Heron Botanicals |
| *Polygonum cuspidatum* | Heron Botanicals<br>(Organic cultivation, China) | Organoleptic, Heron Botanicals | negative testing for pesticides, sulfur dioxide, aerobic plate count and yeast & mold | 30, 60, 90% ETOH by Heron Botanicals |
| *Scutellaria baicalensis* | Heron Botanicals<br>(Organic cultivation, China) | Organoleptic, Heron Botanicals | negative testing for pesticides, sulfur dioxide, aerobic plate count and yeast & mold | 30, 60, 90% ETOH by Heron Botanicals |
| *Cryptolepis sanguinolenta* | Heron Botanicals<br>(Wild harvested, Ghana) | HPTLC, The Institute for Food Safety and Defense<br>Organoleptic, Heron Botanicals | negative testing for Pb, Cd, Hg, As, aerobic plate count and yeast & mold | 30, 60, 90% ETOH by Heron Botanicals |
| *Cistus incanus* | BioPure Healing Products <sup>TM</sup> | DNA Species Identification, NSF International | Negative testing for aerobic plate count, e. coli, coliforms and yeast & mold | 45% ETOH by BioPure Healing Products (aerial parts). DNA analysis reports Cistus Incanus and Cistus albidus are genetically indistinguishable |
| Monolaurin | Lauricidin<sup>TM</sup> |  | N/A |  |
| Colloidal silver | Argentyn 23<sup>TM</sup> |  |  |  |

Botanical medicines were sourced from KW Botanicals (San Anselmo, California) and Heron Botanicals (Kingston, Washington). Botanicals were identified via macroscopic and organoleptic methods and voucher specimens are on file with the respective production facilities. Most botanical medicines were provided as alcohol extracts at 30%, 60%, and 90% alcohol, and the alcohol used was also tested separately as a control in different dilutions. Monolaurin (Lauricidin ™ brand) (Dissolved in 100% DMSO), and colloidal silver (Argentyn ™ brand) were purchased commercially. LL37 and a control was obtained from Taylor Made Pharmacy in Nicholasville, KY. Citrosept ™ (Cintamani, Poland) and Nutribiotic ™ grapefruit seed extract products and a control were purchased commercially. See Table 2 for additional details on sourcing, testing and validation of botanical and natural medicines used.

Doxycycline (Dox) and cefuroxime (CefU) (Sigma-Aldrich, USA) were dissolved in suitable solvents [2, 3] to form 5 mg/ml stock solutions. The antibiotic stocks were filter-sterilized by 0.2 μm filter and stored at −20°C.

### Microscopy

*B. burgdorferi* spirochetes and aggregated microcolonies treated with natural products or control drugs were stained with SYBR Green I and PI (propidium iodide) and checked with BZ-X710 All-in-One fluorescence microscope (KEYENCE, Itasca, IL, USA). The bacterial viability was performed by calculating the ratio of green/red fluorescence to determine the ratio of live and dead cells, as described previously [4, 5]. The residual cell viability reading was obtained by analyzing three representative images of the same bacterial cell suspension taken by fluorescence microscopy. To quantitatively determine the bacterial viability from microscope images, Image Pro-Plus software was employed to evaluate fluorescence intensity as described previously [6].

### Evaluation of natural products for their activity against *B. burgdorferi* stationary phase cultures

*B. burgdorferi* B31 was cultured for 7 days in microaerophilic incubator (33°C, 5% CO_2_) as stationary phase cultures (~10^7-8^ spirochetes/mL). To evaluate potential anti-persister activity of the natural products, their stocks and their control solvents were added to 100 µL of the *B. burgdorferi* stationary phase culture in 96-well plate to obtain the desired concentrations. The botanical medicines and natural product extracts were tested with the concentration of 1%, 0.5% and 0.25% (v/v); antibiotics of daptomycin, doxycycline and cefuroxime were used as control at a final concentration of 5 μg/ml. All the tests mentioned above were run in triplicate. The microtiter plates were sealed and incubated at 33°C without shaking for 7 days with 5% CO_2_.

### Subculture studies to confirm the activity of the top natural product hits

For the subculture study, 1 mL *B. burgdorferi* stationary phase culture was treated by natural products or control drugs in 1.5 ml Eppendorf tubes for 7 days at 33 °C without shaking. Next, cells were centrifuged, and cell pellets were washed with fresh BSK-H medium (1 mL) followed by resuspension in fresh BSK-H medium without antibiotics. Then 50 μl of cell suspension was inoculated into 1 ml of fresh BSK-H medium for subculture at 33 °C, 5% CO_2_. Cell growth was monitored using SYBR Green I/PI assay and fluorescence microscopy after 7-20 days.

## Results

### Evaluation of activity of natural product extracts against stationary phase *B. burgdorferi*

We tested a panel of botanical medicines and natural product extracts and their corresponding controls against a 7-day old *B. burgdorferi* stationary phase culture in 96-well plates incubated for 7 days. Table 1 summarizes the activity of these natural product extracts against the stationary phase *B. burgdorferi* culture at 1%, 0.5% and 0.25%. Among them, 12 natural product extracts at 1% were found to have strong activity against the stationary phase *B. burgdorferi* culture compared to the control antibiotics doxycycline and cefuroxime (Table 1). To eliminate auto-fluorescence background, we checked the ratio of residual live cells and dead cells by examining microscope images as described previously [7]. Using fluorescence microscopy, we confirmed that 1% *Cryptolepis sanguinolenta, Juglans nigra*, and *Polygonum cuspidatum* could eradicate almost all live cells with only dead and aggregated cells left as shown in Figure 1. At 0.5% concentration, 11 natural product extracts (*Polygonum cuspidatum* 60% EE, *Cryptolepis sanguinolenta* 60% EE, *Artemesia annua* 90% EE, *Juglans nigra* 30%-60% EE, *Uncaria tomentosa* WE, *Artemesia annua* 60% EE, *Polygonum cuspidatum* 90% EE, *Scutellaria baicalensis* 30%-90% EE) still exhibited stronger activity than the current clinically used doxycycline and cefuroxime (Table 1; Figure 1). Among them, the most active natural product extracts were *Cryptolepis sanguinolenta* 60% EE, *Polygonum cuspidatum* 60% EE, *Artemesia annua* 90% EE, *Juglans nigra* 60% EE, *Uncaria tomentosa* WE, *Artemesia annua* 60% EE, because of their outstanding activity even at 0.25%, as shown by better activity than control drugs (Table 1 and Figure 1). In particular, 0.25% *Cryptolepis sanguinolenta* could eradicate or dissolve all the *B. burgdorferi* cells including aggregated forms as we found rare live and even dead cells with SYBR Green I/PI microscope observation (Figure 1). Although *Juglans nigra* could eradicate almost all stationary phase *B. burgdorferi* cells at 0.5% (Figure 1), it could not kill the aggregated microcolony form at 0.25% as shown by many live (green) microcolonies by SYBR Green I/PI microscopy. Although the plate reader data showed *Polygonum cuspidatum* 60% ethanol extract had the strongest activity at 0.25%, the microscope result did not confirm it due to higher residual viability than that of *Cryptolepis sanguinolenta* and *Juglans nigra* (Figure 1).

**Figure 1.**
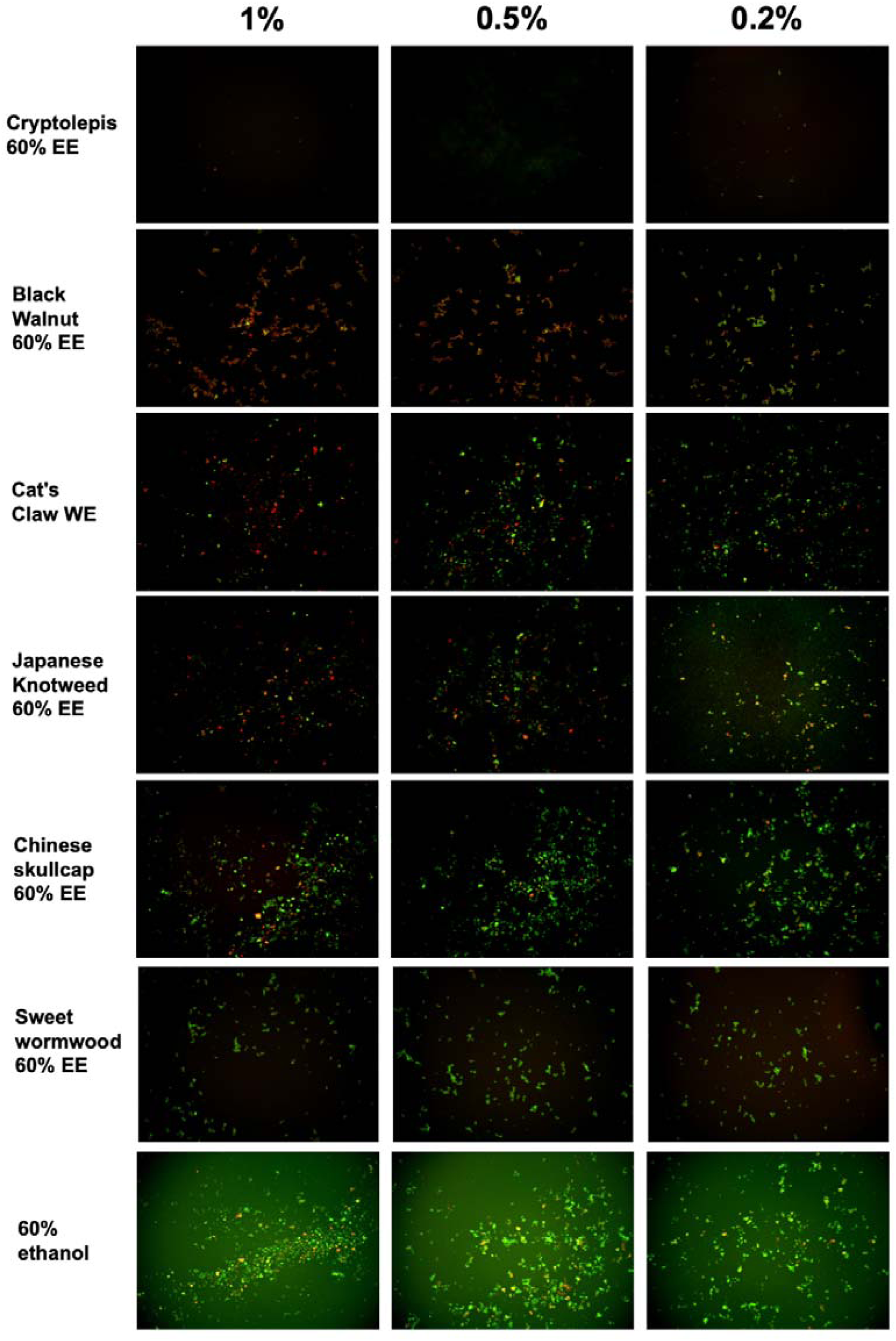
Effect of natural product extracts on the viability of stationary phase *B. burgdorferi*. A 7-day old *B. burgdorferi* stationary phase culture was treated with the natural product extracts at 1%, 0.5% and 0.2% for 7 days followed by staining with SYBR Green I/PI viability assay and fluorescence microscopy.

We also tested several other herbs and substances that are used by Lyme patients including *Stevia rebaudiana, Andrographis paniculata*, Grapefruit seed extract, *Ashwagandha somnifera*, Colloidal silver, Lauricidin, and antimicrobial peptide LL-37, but found they had little or no activity against stationary phase *B. burgdorferi* cells.

### MIC values of the active natural product extracts

Because the activity of antibiotics against non-growing *B. burgdorferi* is not always correlated with their activity against growing bacteria [7], we therefore determined the MICs of these natural product extracts against the replicating *B. burgdorferi* as described previously [8]. The MIC values of some natural product extracts such as *Artemesia annua, Juglans nigra, Uncaria tomentosa* were quite high for growing *B. burgdorferi*, despite their strong activity against the non-growing stationary phase *B. burgdorferi* cells (Table 1). On the other hand, the top two active natural product extracts *Cryptolepis sanguinolenta* and *Polygonum cuspidatum* showed strong activity against the growing *B. burgdorferi* with a low MIC (0.03%-0.06% and 0.25%-0.5% respectively) and also non-growing stationary phase *B. burgdorferi* (Table 1).

### Subculture studies to evaluate the activity of natural product extracts against stationary phase *B. burgdorferi*

To confirm the activity of the natural product extracts in eradicating the stationary phase *B. burgdorferi* cells, we performed subculture studies as previously described [6]. We further tested the top active natural product extracts (*Cryptolepis sanguinolenta, Polygonum cuspidatum, Artemesia annua, Juglans nigra*, and *Scutellaria baicalensis*) to ascertain if they could eradicate stationary phase *B. burgdorferi* cells at 1% or 0.5% by subculture after the treatment (Table 1). Treatment with 1% *Cryptolepis sanguinolenta* extract caused no regrowth in the subculture study (Table 1, Figure 2). However, the other natural product extracts including *Polygonum cuspidatum, Artemesia annua, Juglans nigra*, and *Uncaria tomentosa* could not eradicate *B. burgdorferi* stationary phase cells as many spirochetes were still visible after 21-day subculture (Table 1, Figure 2). At 0.5%, all the natural product extracts treated samples grew back after 21-day subculture (Table 1, Figure 2), however, only one of the three *Cryptolepis sanguinolenta* extract treated samples grew back. This indicates that 0.5% *Cryptolepis sanguinolenta* extract still has strong activity and could almost eradicate the stationary phase *B. burgdorferi* cells. By contrast, the clinically used antibiotics doxycycline and cefuroxime at clinically relevant concentration (5 μg/ml) could not sterilize the *B. burgdorferi* stationary phase culture, since spirochetes were visible after 21-day subculture (Table 1).

**Figure 2.**
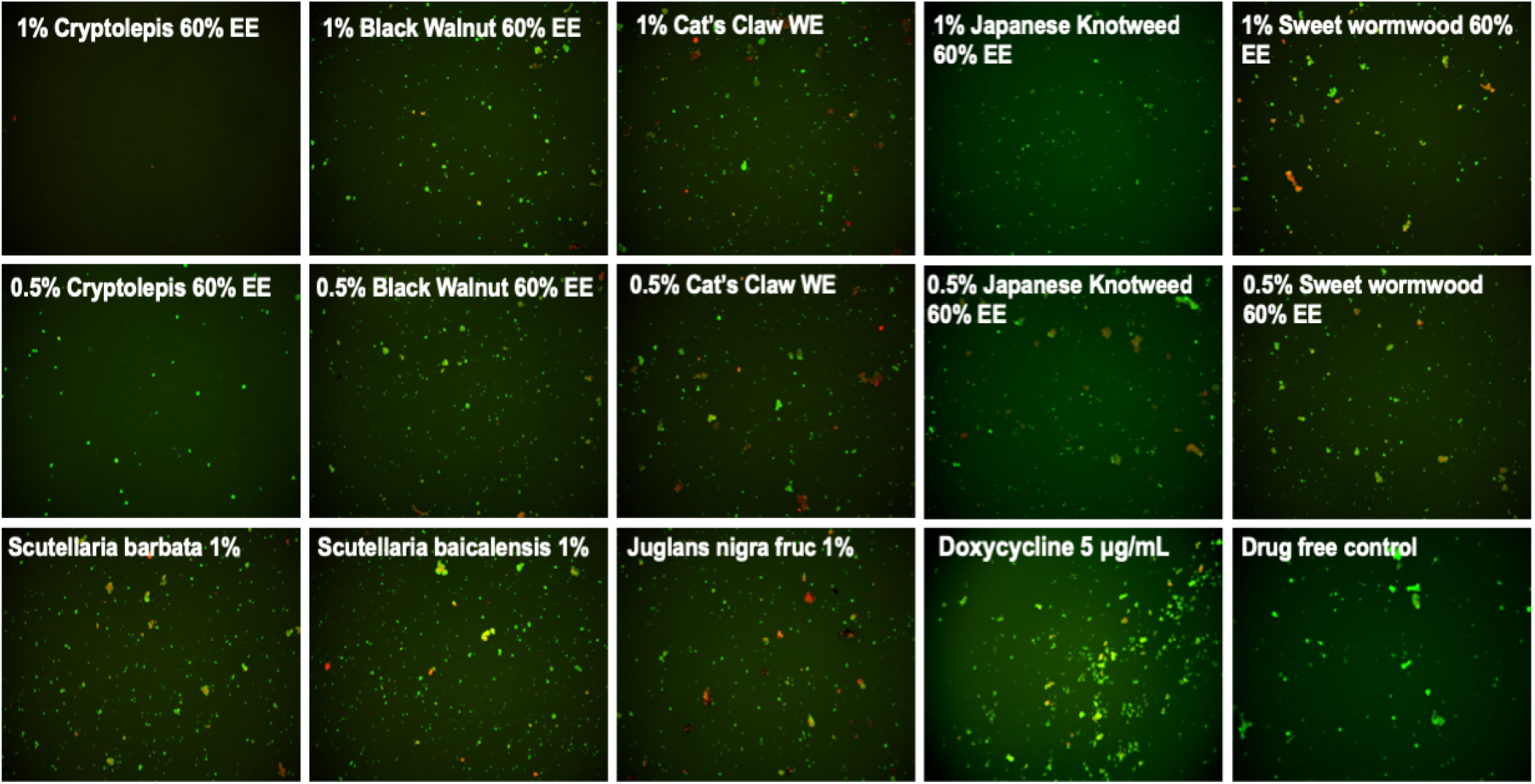
Subculture of *Borrelia burgdorferi* after treatment with natural product extracts. A 7-day stationary phase *B. burgdorferi* culture was treated with the indicated natural product extracts for 7 days followed by washing and resuspension in fresh BSK-H medium and subculture for 21 days. The viability of the subculture was examined by SYBR Green I/PI stain and fluorescence microscopy.

## Discussion

In this study, we evaluated a panel of botanical medicines and natural products commonly used by some patients to manage their persisting symptoms of Lyme disease and found that indeed some of them have strong activity against *B. burgdorferi*. These include *Cryptolepis sanguinolenta, Polygonum cuspidatum, Juglans nigra, Artemisia annua, Uncaria tomentosa, Cistus incanus*, and *Scutellaria baicalensis*. These findings may provide some basis for the clinical improvement of patients who take these medicines and also indirectly suggest their persisting symptoms may be due to persistent bacteria that are not killed by conventional Lyme antibiotic treatment. Since these herbs contain different components and their effects in patients may also be due to their effects on host systems in addition to their potent antimicrobial effect. Surprisingly, *Andrographis paniculata, Stevia rebaudiana (50)*, Colloidal silver (Argentyn 23), Monolaurin (Lauricidin), *Dipsacus spp*, and *Ashwagandha somnifera*, which are assumed or previously reported to have anti-borrelia activity, did not show significant activity against either stationary phase or growing *B. burgdorferi* in our in vitro study, and it is possible that their beneficial effects seen in patients may be in part due to their activity on host immune system.

### Cryptolepis sanguinolenta

*Cryptolepis sanguinolenta* is a plant indigenous to Africa where it has been used in traditional medicine to treat malaria, TB, hepatitis, and septicemia (51). In addition to the various uses documented in ethnomedicine, *Cryptolepis sanguinolenta* has been shown in preclinical studies to have anti-inflammatory (52, 53) antibacterial (54–59), Anti-fungal (55, 60), anti-amoebic (61) and anti-malarial (62–65) properties. Two preliminary clinical studies have shown *Cryptolepis* to have significant efficacy in treating uncomplicated malaria without signs of overt toxicity (66, 67).

While multiple secondary metabolites with antimicrobial activity have been identified, an alkaloid called cryptolepine has been the most well-studied to date. Cryptolepine’s antimicrobial activity is thought to be secondary to multiple mechanisms of action including both bactericidal and bacteriostatic effects (54). More specifically, cryptolepine has been shown to cause morphologic changes and cellular breakdown (60, 68), as well as DNA intercalating and topoisomerase II inhibiting effects (69–73).

It should be noted that, in addition to cryptolepine, other constituents in *Cryptolepis sanguinolenta* have also been shown to have antimicrobial activity (56). A concept in botanical medicine postulates that using a whole plant extract offers several potential advantages over the use of a single constituent, including multiple mechanisms of action, synergism and, in some cases, improved bioavailability as well as less side effects. An example of the clinical benefit in using whole plant extracts over single constituents or analogues may be emerging from the current use of artemisinin-based combination therapy (ACT) for malaria where significant resistance has emerged (74, 75) whereas preliminary studies show improved efficacy and reduce side-effects compared to whole plant treatment (76, 77).

*Cryptolepis sanguinolenta* is generally well tolerated and few side effects have been documented in humans even with its relatively long-term use in parts of China and India. Rat studies indicate that doses of the extract up to 500 mg/kg are relatively safe (78). However, higher doses induced CNS toxicity, thrombocytosis and inflammation in target organs. LD50 was estimated at greater than 3000 mg/kg (79). *Cryptolepis sanguinolenta* was shown in a rat model to lower testosterone levels and reduce sperm counts (80). However, this study was done with a preparation from the leaf of the plant and in Western botanical medicine the root is generally used. Additional studies would be needed to clarify if *Cryptolepis sanguinolenta* has anti-androgenic or anti-spermatogenic effects in humans.

Importantly, a novel finding of this current study is the fact that *Cryptolepis sanguinolenta* has strong activity against growing *B. burgdorferi* with low MIC and also non-growing stationary phase *B. burgdorferi* (Table 1, Fig. 1 and 2). Due to the fact that *Cryptolepis sanguinolenta* is traditionally used against malaria, in the Lyme treatment community it has been used for treatment of *Babesia spp* (81) which can be a co-infecting, malaria like organism. To our knowledge, the anti-*Borrelial* effect of *Cryptolepis sanguinolenta* has not previously been documented and further in vitro and in vivo studies are warranted to investigate the potential role *Cryptolepis sanguinolenta* may serve in the treatment of Lyme disease.

### Juglans nigra

*Juglans nigra* and its constituents have been shown to have anti-oxidant, anti-bacterial, anti-tumor and chemoprotective effects (82–84). Previous in vitro testing has documented that *Juglans nigra* exhibited bacteriostatic activity against log phase spirochetes of *Borrelia burgdorferi* and *Borrelia garinii* and bactericidal activity against *Borrelia* round bodies (85). Two different commercially available botanical formulations which contain *Juglans nigra* were also recently shown to have activity against log phase spirochetes of *B. burgdorferi* strain GCB726, round bodies and biofilm formation in in vitro testing (86). This current study adds to the research on the potential anti *Borrelia* activity of *Juglans nigra* which has been shown to have several constituents (87) with antimicrobial properties including juglone (5-hydroxy-1,4-naphthalenedione), phenolic acids, flavonoids, and catechins (including epigallocatechin gallate) (88–93). Further studies are needed to elucidate which constituents have anti-borrelial activity.

*Juglans nigra* is well tolerated with uncommon side effects. In some individuals, it can cause gastrointestinal disturbance/upset stomach (Natural Medicines Monograph: Black Walnut accessed 3/4/19). There can be some cross reactivity in terms of allergy in those allergic to tree nuts or walnuts, as well as cases of dermatitis reported in humans and laminitis in horses (94–96),. In addition, *Juglans nigra* can induce changes in skin pigmentation (97, 98). The active compound juglone was found to have an oral LD50 in rats of 112 mg/kg (99).

### Polygonum cuspidatum (Japanese Knotweed)

*Polygonum cuspidatum* is commonly used by Lyme disease patients manage their symptoms (81) and its constituents have been shown to have anti-tumor, antimicrobial, anti-inflammatory, neuroprotective, and cardioprotective effects (100–104). One of the active constituents found in *Polygonum cuspidatum* is a polyphenol called resveratrol. Previous in vitro testing has documented that resveratrol exhibited activity against log phase spirochetes of *Borrelia burgdorferi* and *Borrelia garinii*, minimal activity against borrelia round bodies, and no significant activity against borrelia associated biofilms (85). Emodin (6-methyl-1,3,8-trihydroxyanthraquinone), another active constituent in *Polygonum cuspidatum*, has been shown to have activity against stationary phase B. burgdorferi cells (105). Preclinical research has documented *Polygonum cuspidatum* to have antibacterial effects against *Vibrio vulnificus* (106), *Streptococcus mutans* (107) and streptococcus associated biofilms (108). The antibacterial activity of *P. cuspidatum* has been attributed to its stilbenes (including resveratrol) and hydroxyanthraquinone content (109).

*Polygonum cuspidatum* has been found to have minimal toxicity in animal and human studies, although gastrointestinal upset and diarrhea can occur but resolves with decreasing or stopping the intake (110, 111). In safety studies of a purified product, trans-resveratrol did not cause any adverse effects in rats at up to 700 mg/kg bw/day when administered for up to 90 days (112). While few studies have been performed in humans, a 2010 review found that it is well absorbed, rapidly metabolized, mainly into sulfo and glucuronide conjugates which are eliminated in urine. Resveratrol seems to be well tolerated and no marked toxicity was reported. These data are important in the context of human efficacy studies, and they provide further support for the use of resveratrol as a pharmacological drug in human medicine (113). Interestingly, intestinal bacteria played an important role in the metabolism (114).

### Artemisia annua

*Artemisia annua* (Sweet wormwood also called Chinese wormwood and Qing Hao) is a medicinal plant that has been used for medicinal purposes for over 2000 years (115) and the isolation of an active constituent called artemisinin by was awarded the Nobel Prize in 2015 in recognition of artemisinin’s role in significantly reducing the morbidity and mortality associated with malaria (116–118). The anti-*Borrelia* activity of *Artemisia annua* found in this current study adds to the fact that artemisinin has previously been shown to have significant activity against stationary phase *B. burgdorferi* persisters in in vitro models (36, 119). A small pilot study demonstrated that a synthetic analog to artemisinin, called artesunate, showed a significant reduction in short term memory impairment in patients with Lyme disease when combined with intravenous ceftriaxone (120).

Artemisinin’s mechanism of action for treating Plasmodium infections is not completely understood (121), but is thought to be related to its ability to generate free radicals that damage parasite proteins (122, 123).

The artemisinin content of the *Artemisia annua* sample used in the present study was confirmed to be 0.11% by high-performance liquid chromatography/UV-visual spectroscopy at the Institute for Food Safety and Defense (Centralia, WA). Good quality *Artemisia annua* should generally contain >0.3% artemisinin. Despite suboptimal levels of artemisinin present in the *Artemisia annua* used for the present study, both 60% and 90% alcohol extracts of *Artemisia annua* exhibited better activity against stationary phase *B. burgdorferi* compared to the control antibiotics cefuroxime and doxycycline. This is consistent with the previous in vitro data demonstrating artemisinin’s ability to reduce round bodies of *B. burgdorferi* (36).

*Artemisia annua* is generally considered safe provided that the product administered is free of or low in thujone and other terpene derivatives that are potentially neurotoxic (124). Rat studies found that the NOAEL (no-observed-adverse-effect-level) of *Artemisia annua* extract in rats was estimated to be equivalent to 1.27 g/kg/day in males and 2.06 g/kg/day in females) or more (125). In humans, *Artemisia annua* has been used safely in doses up to 2250 mg daily for up to 10 weeks (124), and 1800 mg daily have also been used safely for up to 6 months (124). Some gastrointestinal upset including mild nausea, vomiting (more rare), and abdominal pain can occur at higher doses (126, 127).

### Scutellaria baicalensis

*Scutellaria baicalensis* and its constituents have been shown to have neuroprotective, antioxidant, anti-apoptotic, anti-inflammatory and anti-excitotoxicity (128–131), One of the active constituents found in *Scutellaria baicalensis*, baicalein, was found to exhibit in vitro activity against various morphologic forms of *B. burgdorferi* and *B. garinii*, including log phase spirochetes, latent round bodies and biofilm formations (132). This current study adds to the research on the anti-*Borrelia* activity of *Scutellaria baicalensis*. This botanical and/or baicalein have also been shown to have antimicrobial activity (133, 134), synergistic effects with antibiotics (135–139) and reduce biofilm formation in *Pseudomonas aeruginosa* models (140, 141).

*Scutelaria baicalensis* has been used safely in clinical use (142–144), and has a long historical record of safety. There are reports of sedation and it has been shown to be active on the GABA receptor sites (though this is frequently used to help anxiety and sleep)(145) (145–147). A medical food combination of purified *Scutellaria baicalensis* and the bark of *Acacia catechu* containing baicalin and catechin, concentrated and standardized to greater than 90% purity (Limbrel ™, Move Free Advanced ™) caused reversible liver damage in at least 35 cases, with a calculated estimated incidence of approximately 1 in 10,000 (148, 149). These commercial products have since been withdrawn from the market. Similar hepatotoxicity is generally not seen from the whole plant extract. Despite the case reports of hepatotoxicity, a dose of 1000 mg/kg daily was identified as the no-observed-adverse-effect level (NOAEL) for this commercial product was given in animal studies for 90 days (150). Another study demonstrated no teratogenicity on *Scutelaria baicalensis* when given to pregnant mice at doses up to 32g/kg/day (151).

### Uncaria tomentosa

*Uncaria tomentosa* is an important medicinal plant from South and Central America and has been shown to have neuroprotective effects in preclinical studies (152) (and in preliminary human studies has been shown to improve quality of life in individuals with cancer (153), enhanced DNA repair (154), and symptom improvement in individuals with rheumatoid arthritis (155) and osteoarthritis (156). The potential antimicrobial effects of *Uncaria tomentosa* have not been widely evaluated. In a non-peer reviewed publication, *Uncaria tomentosa* was reported to have anti-borrelial effects in an in vitro model (157). *Uncaria tomentosa* has been shown in peer reviewed research to have antimicrobial effects against human oral pathogens (158, 159).

*Uncaria tomentosa* has been found to be safe and to have minimal side effects in a variety of animal and human studies (154). Human studies ranging from four weeks (156) to 52 weeks (155) demonstrated side effects comparable to placebo. While gastrointestinal complaints such as nausea, diarrhea, abdominal pain, and anemia, were reported, it was thought that the group of solid tumor patients had experienced health issues from disease progression and not necessarily from the *Uncaria* (153). One case report was made of allergic interstitial nephritis in a patient with SLE whose kidney function worsened when taking an *Uncaria tomentosa* product and improved upon discontinuation (160). LD50 of several different preparations of *Uncaria tomentosa* was found to range from 2-8 g/kg bodyweight (McKenna DJ, Jones K, Hughes K, Humphrey S, editors. Botanical Medicines. The desk reference for Major Herbal Supplements. 2nd ed. The Haworth Herbal Press, Binghamton, NY USA 2002). Another study calculated the acute median lethal dose in mice to be greater than 16 g/kg body weight (161).

### Cistus creticus

It has been proposed that *Cistus incanus* and *Cistus creticus* are synonymous (theplantlist.org) while other sources have suggested that *Cistus creticus* is a subspecies of *Cistus incanus* (162). Preliminary clinical studies have shown significant improvement in upper respiratory infection and inflammatory markers in patients taking *Cistus incanus* (163, 164), A volatile oil extract of *Cistus creticus* has been shown to have anti-borrelial effects in an in vitro model (165). Additional in vitro studies have shown *Cistus creticus* to have antimicrobial effects against several bacteria including *Pseudomonas aeruginosa, Klebsiella pneumoniae* (162), *Streptococcus oralis, Staphylococcus aureus, Porphyromonas gingivalis, Prevotella intermedia, Fusobacterium nucleatum* and *Parvimonas micra* (166). *Cistus creticus* also demonstrated significant inhibition of *Streptococcus mutans* biofilm formation (166) and reduction in bacterial adherence to enamel (167). *Cistus creticus* has been shown to contain several active constituents (168), including carvacrol (165). Given that our lab previously documented carvacrol to have a significant activity against log and stationary phase *B. burgdorferi* cells, it is possible that the carvacrol content in the *Cistus incanus* sample tested in the present study contributed to the significant reduction in long and stationary phase *B. burgdorferi* cells in the present study

*Cistus incanus* plant extracts have been used for centuries in traditional medicine without reports of side effects or allergic reactions (169). In a randomised placebo controlled study of 160 patients, 220 mg per day *Cistus incanus* was well tolerated with less adverse effects than in the placebo group (163). In a similar study comparing *Cistus incanus* to green tea, less adverse effects was again seen in the *Cistus incanus* group compared to the green tea group (164). While pharmacokinetic safety data is sparse, a cell culture study showed that *Cistus incanus* did not cause any adverse changes on cell proliferation, survival, or cellular receptor function (169).

### Grapefruit seed extract

Grapefruit seed extract (GSE) was previously reported to have activity against motile and cystic morphologic forms of borrelia bacteria in an in vitro model in a 2007 publication (170). In contrast, the current study did not demonstrate any meaningful activity against log phase or stationary phase *B. burgdorferi*. There are several potential reasons to explain the difference in results between the current study and the 2007 study including differences in GSE formulations and/or different borrelia species used in culture. In the current study we used *B. burgdorferi* strain b31 whereas the 2007 study cites “*B. afzelii* ACA-1 “was used. While both studies used Citrosept ™ brand GSE the formulation has been modified since 2007 and currently holds an “organic” designation. Because previous studies have documented several contaminants in commercial GSE formulations, including Benzalkonium chloride, triclosan and methylparaben (171–173), we screened the GSE products for contaminants prior to inclusion in our present study. The Citrosept ™ sample was found to have no detectable levels of contaminants and therefore was used as the GSE source in the current study. In contrast, a second commercially available brand of GSE (Nutribiotic ™) did test positive for elevated levels of Benzalkonium chloride, which is a known antimicrobial compound (174) and has been implicated in drug-herb interactions causing potential safety concerns for patients taking GSE (175). The 2007 study did not note testing for contaminants so it is possible that the 2007 formulation of Citrosept ™ contained a contaminant that exerted anti-borrelial activity.

### Stevia rebaudiana

*Stevia rebaudiana* was recently reported to have strong anti-borrelia activity (50). However, in our testing, *Stevia rebaudiana* failed to show activity against *B. burgdorferi*. One possibility to explain this discrepancy is that the study that reported *Stevia rebaudiana* having activity against *B. burgdorferi* did not have appropriate alcohol control and that the anti-borrelial effect seen with the *Stevia rebaudiana* alcohol extract may not be due to *Stevia rebaudiana* but due to a non-specific alcohol effect on the *Borrelia* bacteria. Since we obtained *Stevia rebaudiana* preparation from an experienced herbalist who extracted *Stevia rebaudiana* using a known concentration of alcohol, we were able to know the alcohol concentration in the preparation and when we used proper alcohol controls we did not find *Stevia rebaudiana* to have any activity against *B. burgdorferi* (Table 1).

### Andrographis paniculata

*Andrographis paniculata* (Chuan Xin Lian) has been used to treat febrile diseases and infections caused by syphilis, malaria, and worms, and is recommended as anti-spirochetal treatment in the Buhner Lyme disease book (106). However, we found Andrographis failed to show any apparent activity against *B. burgdorferi* in our in vitro testing. It is possible that Andrographis indirectly acts on the host immune system to kill *B. burgdorferi* or have a non-specific host response. Further studies are needed to test the possible effect of Andrographis on the host immune cells.

Other substances or compounds used by Lyme patients such as Colloidal silver, Monolaurin, Grapefruit seed extract, and antimicrobial peptide LL-37 did not exhibit good activity against *B. burgdorferi* in our testing.

## Conclusion

In conclusion, we tested a panel of herbal natural products that are most commonly used by Lyme disease patients for their activity against *B. burgdorferi* and found several to be highly active including *Cryptolepsis sanguinolenta, Juglans nigra, Polygonum cuspidatum, Uncaria tomentosa, Artemisia annua, Cistus creticus*, and *Scutellaria baicalensis*. However, we found that *Stevia rebaudiana, Andrographis paniculata*, Grapefruit seed extract, colloidal silver, monolaurin, and antimicrobial peptide LL37 had little or no activity against *B. burgdorferi* in our in vitro model. Future studies are needed to identify the active ingredients of the effective herbs and to evaluate their potential for more effective treatment of persistent Lyme disease in animal models and in patients.

While this current study has identified novel new botanical and natural medicines with in vitro anti-*Borrelia* activity, it is also notable that many compounds tested did not show direct anti-*Borrelia* activity despite the fact that they are widely used, with reported clinical efficacy, by patients and practitioners in the community setting. It is important to consider the potential limitations of the in vitro model given that it exists outside of the biological organism. The in vitro model can provide information with regards to direct antimicrobial activity, and while botanical and natural medicines can be effective from direct antimicrobial activity, frequently part of their function is via diverse pathways which are not directly antimicrobial. For example, they can exert effects via anti-inflammatory/anti-cytokine activity, immune system regulation/augmentation, adaptogenic stimulation of cellular and organismal defense systems, and biofilm disruption to name a few (see discussion section). In these activities, the mechanisms of the medicines rely on complex interplay and interaction between different body systems, which can only occur within the intact, living organism. Because the in vitro model is unable to provide information with regards to alternative pathways through which natural botanical medicines act, it is important that future in vivo studies be performed to investigate the activity and efficacy of these and other botanical and natural medicines against Borrelia and other tick-borne diseases.

These types of studies will be of vital importance given the multiple factors at play with the current epidemic of tick-borne diseases in our society and globally. While research is beginning to provide information on novel antibiotic combinations as well as agents previously not used for this purpose (34) that might be effective against the multiple forms of the Borrelia bacteria, there is ongoing concern and care is required regarding issues of responsible stewardship of antibiotic use and antibiotic resistance. It is also important to recognize that, while being cognizant of specific side effects and interactions, botanical and natural medicines generally have a favorable safety profile compared to prescription antibiotics and have a broader spectrum of action with multiple synergistic compounds present within a single plant. Furthermore, using multiple botanical medicines in combination can further increase synergy and lower the risk of pathogen resistance development.

Finally, given the need for novel antimicrobials that are active against the persistent form of the Borrelia bacteria which is difficult to treat even with conventional antibiotic approaches, additional research is critical to identify the active components of the effective hits and evaluate the activity of active botanical medicines in combination against Borrelia persisters in vitro and in vivo in the mouse model of Borrelia infection and in subsequent clinical studies.

## Acknowledgments

We thank herbalists Eric Yarnell, Brian Kie Weissbuch, and Mischa Grieder ND for providing herbal extracts for evaluation in this study and for helpful discussions. We acknowledge the support of this work by the Bay Area Lyme Foundation.

